# A random priming amplification method for whole genome sequencing of SARS-CoV-2 and H1N1 influenza A virus

**DOI:** 10.1101/2021.06.25.449750

**Authors:** Klaudia Chrzastek, Chandana Tennakoon, Dagmara Bialy, Graham Freimanis, John Flannery, Holly Shelton

**Affiliations:** The Pirbright Institute, Pirbright, Woking, UK

**Keywords:** whole genome sequencing, SISPA, SARS-CoV-2, coronavirus, influenza virus, metagenomics, RNA

## Abstract

**Background:** Non-targeted whole genome sequencing is a powerful tool to comprehensively identify constituents of microbial communities in a sample. There is no need to direct the analysis to any identification before sequencing which can decrease the introduction of bias and false negatives results. It also allows the assessment of genetic aberrations in the genome (e.g., single nucleotide variants, deletions, insertions and copy number variants) including in noncoding protein regions.

**Methods:** The performance of four different random priming amplification methods to recover RNA viral genetic material of SARS-CoV-2 were compared in this study. In method 1 (H-P) the reverse transcriptase (RT) step was performed with random hexamers whereas in methods 2-4 RT incorporating an octamer primer with a known tag. In methods 1 and 2 (K-P) sequencing was applied on material derived from the RT-PCR step, whereas in methods 3 (SISPA) and 4 (S-P) an additional amplification was incorporated before sequencing.

**Results:** The SISPA method was the most effective and efficient method for non-targeted/random priming whole genome sequencing of COVID that we tested. The SISPA method described in this study allowed for whole genome assembly of SARS-CoV-2 and influenza A(H1N1)pdm09 in mixed samples. We determined the limit of detection and characterization of SARS-CoV-2 virus which was 10^3^ pfu/ml (Ct, 22.4) for whole genome assembly and 10^1^ pfu/ml (Ct, 30) for metagenomics detection.

**Conclusions:** The SISPA method is predominantly useful for obtaining genome sequences from RNA viruses or investigating complex clinical samples as no prior sequence information is needed. It might be applied to monitor genomic virus changes, virus evolution and can be used for fast metagenomics detection or to assess the general picture of different pathogens within the sample.

## Background

Advances in next-generation sequencing (NGS), including targeted or non-targeted whole genome sequencing (WGS) and computational analyses capable of efficiently processing large amounts of data have enabled us to comprehensively study viral genomes in research and clinical settings. NGS and bioinformatics approaches have been used to identify the causative agents of the outbreaks, outbreak origins, track transmissions or investigate epidemic dynamics, including outbreaks of Ebola, yellow fever and Zika virus (ZIKV) and recently SARS-CoV-2 pandemic [1–12]. For instance, Faria et al. [4], applied WGS to early samples collected from ZIKV infected patient and estimated the date of first introduction of ZIKV into the Americas a year before the first detections in Brazil in 2014. Whole genome sequencing was also successfully applied to recover the HIV-1 genome from the individual known as ‘Patient 0’ and other samples from 1970s to further understand the emergence of HIV-1 in the USA [13]. In early 2020, the first recovered SARS-CoV-2 genomes from eight patients in Wuhan demonstrated an identity of 99.98% [9]. The high level of shared genomic similarity between early SARS-CoV-2 viral genomes clearly suggested that the virus had not been circulating long in the human population but was likely a spill over of an animal coronavirus into humans [9].

WGS is also a powerful tool for the screening of virus evolution including drug resistance mutations/genes, vaccine escape variants, recombination or reassortment, and virulence and pathogenicity factors [14–18]. Recently, Kemp et al. [19] showed intra-patient SARS-CoV-2 virus genetic diversity increased following plasma treatment of a patient with high viral loads. Furthermore, a high rate of mutation in the SARS-CoV-2 virus genome was seen in immunodeficient patients chronically infected with the virus [20].

Since the emergence of SARS-CoV-2 in 2019, and further classification of the initial outbreak as a global pandemic on March 11, 2020, a total of 145M positive cases were registered including 3.07M deaths globally by April 23rd, 2021 (ECDC, 2021). Over that period, there has been an unparalleled global effort to characterise the virus biology and identify genomic changes in the SARS-CoV-2 virus genome that would not be possible without applying whole genome sequencing on collected samples both in research and clinical settings. WGS allowed researchers to identify a novel variant of coronavirus, especially mutations or deletions in the gene encoding the receptor spike protein of the virus to which the predominant protective immunological response is directed [21–26]. Korber et al. [22] performed genomic analysis on 28,576 sequences available in GISAID database (by May 29th, 2020) and shown that a SARS-CoV-2 variant carrying the spike protein amino acid change D614G had replaced the original Wuhan form of the virus across the globe by June 2020. A variant linked to infection among farmed mink SARS-CoV-2 strain, which had the spike protein change Y453F, and subsequently transmitted to humans, was identified in North Jutland, Denmark was reported in late 2020 [27, 28]. In December 2020, The United Kingdom identified a large proportion of SARS-CoV-2 cases that belong to a new single phylogenetic cluster, the B.1.1.7 lineage (alpha variant) following an unexpected rise in cases in South East England, which rapidly became the predominant strain circulating in humans because of increased transmissibility [25, 26, 29, 30]. Furthermore, South Africa also reported a SARS-CoV-2 variant (B.1.351 lineage, beta variant) that had the N501Y mutation like alpha B.1.17 variant but was phylogenetically different. Most recently the delta variant (B. 1.617.2) has been identified with a different sequence and increased transmissibility characteristics [31]. The dynamic changes in SARS-CoV-2 virus genome that have occurred over the period of pandemic, paired with the recent implementation of vaccination programs on a global scale that might further impact variant generation suggests that routine whole genome sequencing of coronavirus genome could be implemented as a vital part of ongoing disease control.

Although WGS has great potential in outbreak tracing and virus monitoring, it is not efficient when there is a low abundance of viral sequence in a sample. In such cases sequence targeted amplicon-based approaches are required to characterise the genome. For example, in diagnostics of clinical sample scenarios where the pathogen is unknown cannot be performed, the often-low abundance of virus genome precludes the opportunity for WGS. Furthermore, to characterise potential mixed viral samples, targeted sets of primers from multiple viruses, would need be required and these would need to be updated frequently since changes in viral genome could influence the targeted primer efficiency. Previous work has shown that a Sequence-Independent, Single-Primer-Amplification (SISPA) technique in combination with Illumina sequencing can be used to recover genetic sequences of negative- and positive-sense single-stranded avian RNA viral sequences [32] and allows the examination of the entire genetic content of a sample, instead of just one particular gene region.

In this study, the performance of four different random priming amplification methods (Figure 1) to recover the sequence of viral RNA from SARS-CoV-2 containing material were compared. We compared the limit of detection necessary for virus identification and metagenomics approaches for each method. In addition we tested the methods on a mixed virus sample containing both SARS-Cov-2 and influenza A(H1N1)pdm09 virus to determine the efficiency in a mixed pathogen scenario which is likely to be seen in the coming Northern Hemisphere winter season.

**Figure 1.**
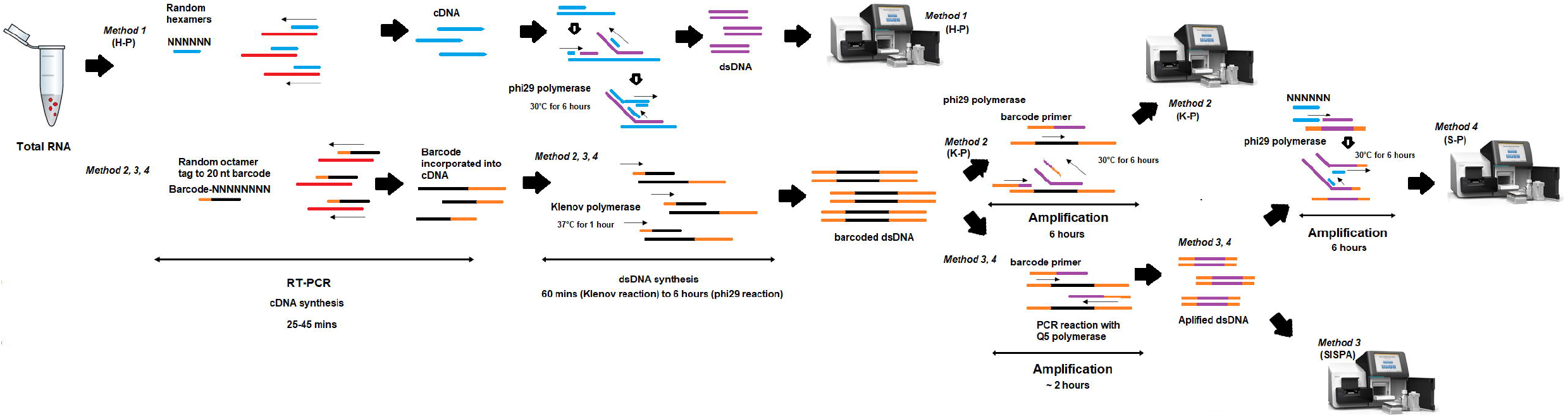
Schema of random amplification methods for whole genome assembly of SARS-Cov-2 virus genome. Method 1 (H-P) is based on the RT-PCR step with random hexamers primer (6Ns) followed by phi29 polymerase isothermal amplification in the presence of 6Ns primer and then library preparation for Illumina sequencing. Method 2 (K-P), random octamer tagged with 20 nucleotide known tag sequence (5’-GACCATCTAGCGACCTCCACNNNNNNNN-3’) (K-8N) was used for RT-PCR step, followed by phi29 polymerase isothermal amplification in the presence of tagged primer K-8N and then library preparation and Illumina sequencing. Method 3, Sequence-Independent, Single-Primer Amplification (SISPA) technique, followed by library preparation and Illumina sequencing. Method 4 (S-P), following SISPA amplification (Method 3), phi29 polymerase isothermal amplification in the presence of random hexamers (6Ns) was applied and then used for Illumina sequencing.

## Material and methods

### Viruses and cells

SARS-CoV-2 virus strains used include hCov-19/England/02/2020 (Eng-2) (EPI_ISL_407073) provided by Public Health England (PHE) and hCov-19/Scotland/EDB1827/2020 (EDB-2) (EPI_ISL_433147), hCov-19/Scotland/EDB2398/2020 (EDB-8) (EPI_ISL_439199), hCov-19/Scotland/EDB2057/2020 (EDB-10) (EPI_ISL_433169) and hCov-19/Scotland/EDB2405/2020 (EDB-12) (EPI_ISL_433169) provided by Dr Christine Tait-Burkard at The Roslin Institute. SARS-CoV-2, hCov-19/England/2/2020 was propagated in Vero E6 cells (ATCC^®^ CRL-1586^™^) and Edinburgh SARS-CoV-2 isolates were propagated in Caco-2 cells (ATCC^®^ HTB-37^™^). Influenza A virus used was A/England/195/09 H1N1 (A(H1N1)pdm09) propagated in the allantoic cavity of 10 day embryonating specific pathogen free hen eggs (VALO Gmb).

Vero E6 cells, Caco-2 cells and MDCK cells were grown in Complete Media, (DMEM (Gibco-Invitrogen, Inc.) supplemented with 10% (v/v) foetal bovine serum (FBS) (Biosera, Inc.) and 1% (v/v) penicillin/streptomycin (Sigma-Aldrich) and maintained at 37°C in 5% CO_2_.

### Infectious virus quantification

Using plaque assay, SARS-CoV-2 was quantified in Vero E6 cells and A(H1N1)pdm09 (IAV) in MDCK cells and expressed as plaque forming units (pfu)/ ml. For SARS-CoV-2, Vero E6 cells were inoculated with 10-fold dilutions of SARS-CoV-2 for 1 h and overlaid with 0.8% (w/v) Avicel medium (1 x MEM Temin’s modification (Gibco), 0.8% (w/v) Avicel^®^ Microcrystalline Cellulose and Sodium Carboxymethylcellulose (FMC BioPolymer), 2% FBS (v/v) (Gibco). After 72 hours of incubation at 37°C, cells were fixed with formalin (VWR) and viral plaques were visualised using 0.1% (w/v) toluidine blue staining (ThermoFisher).

For A(H1N1)pdm09 IAV, MDCKs were inoculated with 10-fold serially diluted samples and overlaid with 0.6% (w/v) agarose (Oxoid) in supplemented DMEM (1× MEM, 0.21% (v/v) BSA V, 1 mM L-Glutamate, 0.15% (v/v) Sodium Bicarbonate, 10 mM Hepes, 1× Penicillin/Streptomycin (all Gibco) and 0.01% (w/v) Dextran DEAE (Sigma-Aldrich, Inc.), with 2 μg/ml TPCK trypsin (SIGMA). They were then incubated at 37°C for 72 hours. Plaques were developed using crystal violet stain containing methanol. RNA extraction

10-fold dilution series for hCov-19/England/02/2020 (Eng-2), starting from 2 x 10^6^ pfu/ ml and a single concentration of A(H1N1)pdm09 IAV (7.4 x 10^6^ pfu/ ml) were RNA extracted from each concentration, using the QIAmp viral RNA minikit (Qiagen) according to manufacturer instructions.

### Quantitative real-time RT-PCR (qRT-PCR)

The viral RNA obtained from each 10-fold dilution of SARS-CoV-2 was titrated using a quantitative real time RT-PCR (qRT-PCR that targeted E-gene of virus genome). In addition, each 10-fold dilution of SARS-CoV-2 virus spiked with a known titre of H1N1 viral RNA which was also quantified using qRT-PCR. The E-Sarbeco assay described by Corman et al. [33] was performed using the Express One-Step Superscript qRT-PCR kit (LifeTechnologies, Paisley, UK). Primers, E_Sarbeco_F (5’-ACAGGTACGTTAATAGTTAATAGCGT-3’) and E_Sarbeco_R (5’-ATATTGCAGCAGTACGCACACA-3’) were used with probe E_Sarbeco_P1 (FAM-ACACTAGCCATCCTTACTGCGCTTCG-BBQ) [33]. For each assay, 15□μl of one-step reaction mix was prepared using 1 × reaction mix, 400□nM forward and reverse primers, 200□nM probe, 0.4□μl Rox, and 2□μl of enzyme. Five microlitres RNA was used per well in a final volume of 20□μl. Cycling conditions were as follows: reverse transcription at 50□°C for 15□min and 95□°C for 20□s, and then 40 cycles of PCR, with each cycle consisting of 95□°C for 3□s and 60□°C for 30□s. RT-qPCR was performed on an Applied Biosystems 7500 Fast real-time PCR instrument (LifeTechnologies). For A(H1N1)pdm09 influenza virus, quantitative analyses of matrix (M) were performed with primers, IAV_F (5’-AGA TGA GTC TTC TAA CCG AGG TCG-3’) and IAV_R (5’-TGC AAA AAC ATC TTC AAG TCT CTG-3’) with IAV_probe (FAM-5’ TCA GGC CCC CTC AAA GCC GA -TAMRA-3’). Briefly, 25□μl of one-step reaction mix was prepared using 1 × reaction mix, 900□nM forward and reverse primers, 100□nM probe, 50 nM Rox, and 0.5 μl SuperScript III One-Step enzyme (ThermoFisher). Two microlitres RNA was used per well in a final volume of 25□μl. T7 RNA transcripts of the M gene with a known concentration was used for the standard curve. Cycling conditions were as follows: reverse transcription at 50□°C for 5□min and 95□°C for 2□mins, and then 40 cycles of PCR, with each cycle consisting of 95□°C for 3□s and 60□°C for 30□s. The qRT-PCR was performed using ABI 7500 FAST machine (LifeTechnologies).

### Random priming (RP)-mediated amplification methods

In this study, four amplification methods were used to achieve a non-selective amplification and recovery of RNA genetic sequences (Figure 1).

Method 1 – Hexamer priming (H-P) uses random hexamer primers (6Ns) for the RT-PCR step with followed by phi29 polymerase (Cytiva illustra^™^), isothermal amplification in the presence of the 6Ns primers according to manufacturer’s instruction to dsDNA for library preparation and Illumina sequencing.

Method 2 – Octamer priming (K-P), uses random octamer primers tagged with a 20-nucleotide tag sequence (5’-GACCATCTAGCGACCTCCACNNNNNNNN-3’) (K-8N) for RT-PCR step, followed by phi29 polymerase isothermal amplification in the presence of tagged primer K-8N to obtain dsDNA necessary for library preparation and Illumina sequencing (Figure 1).

Method 3-Sequence-Independent, Single-Primer Amplification (SISPA) (described previously by Chrzastek et al, 2017) was used to amplified extracted RNA, followed by library preparation and Illumina sequencing.

Method 4-SISPA & phi29 amplification (S-P). Following the SISPA amplification (method 3) and additional phi29 polymerase isothermal amplification in the presence of random hexamers (6Ns) was applied to dsDNA, followed by library preparation and Illumina sequencing (Figure 1).

### RT-PCR

Method 1 (H-P): Random hexamers (50uM, final concentration 2.5uM per reaction) were added to the RT-PCR reaction to synthetize first-stranded cDNA in a 20 μl reaction mixture with 5 μl of viral nucleic acids from each sample, SuperScript IV Reverse Transcriptase (ThermoFisher scientific), and dNTPs (10 μM) according to manufacturer’s instruction. Annealed RNA and RT mix reaction was incubated at 55°C for 10 mins, followed by enzyme inactivation at 80°C for 10 mins.

Method 2 (K-P), Method 3 (SISPA) and Method 4 (S-P): First-stranded cDNA was synthetized in a 20 μl reaction mixture with 5 μl of viral nucleic acids from each sample, 100 μM of primer K-8N (1 μl per reaction), SuperScript IV Reverse Transcriptase (ThermoFisher scientific), and dNTPs (10 μM) following the manufacturer’s instructions. Annealed RNA and RT mix reaction was incubated at 55°C for 10 mins, followed by enzyme inactivation at 80°C for 10 mins.

#### dsDNA synthesis after RT-PCR step

Method 1 (H-P): Genomiphi^™^ V2 DNA Amplification Kit (Cytiva illustra^™^, formerly GE Healthcare Life Sciences) was used for whole genome amplification according to manufacture instruction. Briefly, 5 μl (150-230ng of ssDNA) obtained after RT-PCR reaction was mixed with 5 μl of sample buffer and heated at 95°C for 3mins and then cooled on ice for 4 mins. Subsequently 5 μl of reaction buffer and 0.2 μl of enzyme was added to the reaction mix and incubated at 30°C for 6 hours.

Method 2 (K-P), Method 3 (SISPA) and Method 4 (S-P): After RT-PCR, 20 μl of first-stranded cDNA was heated at 94°C for 3mins and then cooled on ice for 3 mins in the presence of 10uM of primer K-8N (0.5μL per reaction), 10 μM dNTPs (0.5 μl per reaction) in 1× Klenow reaction buffer (NEB). Next, 1 μl of Klenow fragment (NEB) was added to the reaction and incubated at 37°C for 60 min. Following Klenow reaction, dsDNA was cleaned using Agencourt AMPure XP beads (Beckman Coulter) in a ration 1:1. The purified dsDNA was subsequently used as a template for isothermal amplification (Method 2) or PCR amplification (Method 3 and Method 4).

#### Isothermal amplification with phi29 polymerase

Method 2 (K-P): (Figure 1). Whole genome amplification used theGenomiphi^™^ V2 DNA Amplification Kit (Cytiva illustra^™^), like method 1 (H-P). However, for method 2 we modified manufacturers protocol by adding primer K-8N to the reaction. Briefly, 5 μl (5-10 ng) of cleaned dsDNA was added to 4 μl of reaction buffer and 1 μl of 10 μM primer K-8N. Reaction mixture was heated at 95°C for 3mins and then cooled on ice. Next, 5 μl of reaction buffer and 0.2 μl of enzyme was added to the reaction mix and incubated at 30°C for 6 hours follow by enzyme inactivation at 65°C for 10 mins. Following phi29 reaction, dsDNA was cleaned using Agencourt AMPure XP beads (Beckman Coulter) at a ratio 1:1. For quantification of the dsDNA, the Qubit dsDNA HS assay (Invitrogen) was performed according to the manufacturer’s instruction. The purified dsDNA was subsequently used for genome sequencing.

#### PCR amplification

Method 3 (SISPA) and Method 4 (S-P) (Figure 1): Sequence-independent PCR amplification was conducted with 5 μl of purified dsDNA obtained after Klenow reaction in 50 μl of final reaction which contained 1x Q5 High-Fidelity Master Mix (NEB), 2.5 μl of 10 μM primer K (5’-GACCATCTAGCGACCTCCAC-3’) and Nuclease-Free water. The PCR cycling conditions were as follow: 98 °C for 30 s, followed by 35 cycles of 98 °C for 10 s, 55 °C for 30 s, and 72 °C for 1 min, with a final extension at 72 °C for 10 min PCR products were purified using Agencourt AMPure XP beads (Beckman Coulter) ratio 0.6×. For quantification of the ds cDNA, the Qubit dsDNA HS assay (Invitrogen) was performed according to the manufacturer’s instruction. The purified dsDNA was subsequently used for genome sequencing (Method 3, SISPA) or for phi29 isothermal amplification (Method 4).

For method 4, following PCR clean up step, Genomiphi^™^ V2 DNA Amplification Kit (Cytiva illustra^™^, formerly GE Healthcare Life Sciences) was used for whole genome amplification according to manufacture instruction without any modification. The purified dsDNA was subsequently used for Nextera XT libraries preparation (Illumina) (Figure 1).

#### Genome sequencing

A total of 1□ng of dsDNA was used to prepare sequencing libraries using the Nextera XT DNA kit (Illumina). Libraries were analysed on a High Sensitivity DNA Chip on the Bioanalyzer (Agilent Technologies). Pooled libraries were sequenced on a 2×300cycle MiSeq Reagent Kit v2 (Illumina, USA) over two separate Illumina MiSeq runs. The first Miseq run consisted of 10 samples, whereas 38 samples were multiplexed on the second run.

#### Sequence analysis

The quality of sequencing reads was assessed using FastQC ver. 0.11.5 [34]. The reads were quality trimmed with using a quality score of 30 or more, in addition to low-quality ends trimming and adapter removal using Trim Galore ver.0.5.0 (https://github.com/FelixKrueger/TrimGalore). *De novo* assembly was performed using SPAdes de novo assembler (version 3.10.1) (k-mer 33, 55, and 77). Resulting contigs were quality assessed using QUAST (version 5.0.2) [35, 36]. Reference-based orientation and scaffolding of the contigs produced by the assembler were performed using Scaffold_builder version 2.2 [37]. Consensus sequences were re-called based on BWA-MEM mapping of trimmed (but un-normalized) read data to the genome scaffold and parsing of the mpileup alignment. Assembly of reference genomes was performed using BWA-MEM ver. 0.7.17 [38] and Geneious 9.1.2 (https://www.geneious.com). This final consensus sequence representative of the major strain in the viral population was used as a reference genome. Cleaned datasets were mapped against the reference followed by variant calling with LoFreq ver 3.0 [39] to identify the presence of variants arising from inter- or intra-population quasispecies at 3% frequency. Filtering the reads against host genome (Gallus gallus 4.0) was performed using BWA-MEM [38].

#### Metagenomics detection

Three independent methods were used to detect the presence of the viruses in the samples (Figure 5).

##### (1) Assembly

The first method used the contigs assembled by SPAdes assembler using inhouse pipeline. If a contig was larger than 150 bases (i.e., the average size of read) a random 100 bp segment of that contig was sampled. These samples were aligned with BLAST to the nt-database. If any of the sampled reads mapped to a virus, its top ten hits were examined, and the contig it was derived from was aligned to the nt-database with BLAST (allowing a maximum of 10 hits per contig). The resulting BLAST alignments were collated to generate a coverage graph of the contigs along the viruses they mapped to.

##### (2) K-mer analysis

The second method analysed k-mers in individual reads (Figure 5). Each read was inspected using Kraken and its minikraken database to build a report containing the possible organisms the sequences originated from and the number of reads supporting their presence. References for any organisms with a minimum of 100 reads were downloaded and reads were mapped to these references using BWA-mem.

##### (3) Mapping

The final method is the alignment of reads to reference SARS-CoV-19 and A influenza genomes. These alignments were used to generate read depth graphs (Figure 5).

The first assembly method can identify organisms if they are present in the sequencing data in a sufficiently high concentration to be assembled. The second method can detect viruses at a lower concentration. The final method would be sensitive if the references were close to the isolates in the samples. We mark a virus to be present in the sample if there is non-random coverage (e.g., uniform overage, long stretch with coverage) of a closely related viral genome in the plots.

## Results

### Four random amplification methods coupled with Illumina sequencing

In this study, four random-amplification methods coupled with Illumina sequencing were compared for the ability to obtain full genome sequences of SARS-CoV-2 virus (Figure 1). Whole genome amplification (WGA) of RNA material, starts with RNA extraction, followed by conversion of RNA into cDNA and then dsDNA synthesis. Once dsDNA is synthetised can be used directly for library preparation using the Nextera XT DNA (Illumina) or further amplified in PCR or isothermal reactions before being used for library preparation. To produce method 1 (H-P), dsDNA following a RT-PCR step with SuperScript^™^ IV One-Step RT-PCR system (ThermoFisher) with random-hexamer primers, a simple, isothermal random-hexamer-primed, phi29 DNA polymerase-based whole genome amplification was applied. For Method 2 (K-P), Method 3 (SISPA), Method 4 (S-P), in RT-PCR step the hexamer primer was replaced with primer K-8N (Material and Method section). This primer (K-8N) contains a known tag (called here “K”) that is linked to the random octamer (8-N).

Following the RT-PCR step the tag is incorporated randomly into cDNA. Klenow DNA polymerase was used generate dsDNA in an isothermal reaction (Material and Methods section). The final product obtained after RT-PCR and Klenow reactions in Methods 2, 3 and 4 is tagged dsDNA (“K” sequence incorporated into dsDNA). The dsDNA obtained was then used for isothermal (Method 2), or PCR-based amplification (Method 3 and 4). In method 2, the focus was to use an isothermal reaction for amplification and elongate the dsDNA fragments. For that reason, we used multiple displacement amplification (MDA) by Φ29 DNA polymerase and a mix of hexamer and K-8N primer. Finally, for methods 3 (SISPA) and method 4 (S-P) PCR-based amplification was used, where the aim was to amplify dsDNA using primer K (Material and Methods section) that binds to the primer tag so that the tag works as a primer binding extension site in PCR reaction. Method 4 (S-P) had an additional MDA step after PCR to amplify and elongate the template by Φ29 DNA polymerase using only the hexamer primers without any modification of the protocol (Genomiphi^™^ V2, Material and Methods section).

### Comparison of the methods to sequence the whole genome of SARS-CoV-2 when abundant genetic material was present

In this study assembly of full or near full genome (≥ 97% genome coverage) of SARS-CoV-2 virus was achieved using all four amplification methods tested when a high titre of ENG-2 virus was analysed (2.6×106 pfu/ml, CT value: 12.22) (Table 1). Under the conditions of abundant genetic material, the SISPA method (Method 3), produced the highest number of reads that mapped to the SARS-CoV-2 reference genome and the highest average depth of genome coverage (Table 1, Suppl. Figure 1). The percentage of reads mapped to the reference SARS-CoV-2 virus genome was 47.35% and 14.79% for SISPA (method 3) and S-P (method 4), respectively whilst for H-P (method 1) and K-P (method 2) amplification was below 1% of total sequencing reads generated (Table 1). The average coverage depth at this concentration of SARS-CoV-2 virus was 13486.11 (SD=15324.3) for SISPA (method 3) versus 835.44 (SD=1333.5) for S-P (method 4), followed by 72.83 (SD=68.406) for K-P (method 2 and 29.10 (SD=23.8) for H-P (method 1) (Table 1). Detailed statistics for SARS-CoV-2 viral genome assembly using SISPA (method 3) is shown in Table 2 and demonstrates that at the high virus titre, both reference mapping and de novo assemblies produced full genome sequence with high depth of coverage per gene. Depth of coverage being above 10,000 nucleotides per base for the viral genes; orf1ab, orf7b, orf8, N, orf10 genes and above 2,000 nucleotides per base for S, orf3a, E, M, orf6 (Table 2 and Suppl. Figure 2).

**Table 1.** The comparison of performance of four different random priming amplification methods to recover RNA viral genetic material of SARS-CoV-2 genome. SARS-CoV-2 was quantified by plaque assay titration on Vero E6 cells (pfu/ml) and qRT-PCR (CT value). For metagenomics, three independent methods were used to detect the presence of the virus in the samples. Kraken, each read was inspected using Kraken and its database to build a report containing the possible organisms the sequences originated from and the number of reads supporting their presence. Blast, if the contig assembled by SPAdes using inhouse pipeline was larger than 150 bases, a random 100 bp segment of that contig was sampled. These samples were aligned with BLAST to the nt-database. The final method, Align is the alignment of sequencing reads to customised reference database.

| Amplication method | Virus titre (pfu/mL) | CT value | Total number of sequencing reads | BWA*-MEM aligned to SARS-CoV-2 genome | The percentage of reads mapped to SARS-CoV-2 out | The percentage of SARS-CoV-2 genome covered | The average depth of coverage per base | Standard deviation (SD) | The number of unmapped reads to SARS-CoV-2 virus genome | Kraken | Blast | Align |
| --- | --- | --- | --- | --- | --- | --- | --- | --- | --- | --- | --- | --- |
| 1 | 2.6x10<sup>6</sup> | 12.22 | 1399928 | 4502 | 0.32 | 97.68 | 29.10 | 23.8 | 1395426 | N | Y | Y |
| H-P | 2.6x10<sup>5</sup> | 15.99 | 1120143 | 1757 | 0.16 | 93.99 | 10.22 | 12.76 | 1118386 | N | Y | Y |
|  | 2.6x10<sup>4</sup> | 18.99 | 1535429 | 1729 | 0.11 | 72.79 | 8.62 | 24.28 | 1533700 | N | Y | Y |
|  | 2.6x10<sup>3</sup> | 22.4 | 635446 | 142 | 0.02 | 27.22 | 0.74 | 1.66 | 635304 | N | N | N |
|  | 2.6x10<sup>2</sup> | 25.34 | 1100547 | 794 | 0.07 | 48.81 | 4.82 | 8.91 | 1099753 | N | N | N |
|  | 26 | 29.34 | 966273 | 45 | 0 | 10.22 | 0.27 | 0.97 | 966228 | N | N | N |
|  | 2.6 | 32.36 | 973044 | 3 | 0 | 0.75 | 0.01 | 0.14 | 973041 | N | N | N |
|  | 0 | 35.04 | 739156 | 1 | 0 | 0.12 | 0.00 | 0.03 | 739155 | N | N | N |
| 2 | 2.6x10<sup>6</sup> | 12.22 | 1414247 | 10798 | 0.76 | 99.36 | 72.83 | 68.41 | 1403449 | Y | Y | Y |
| K-P | 2.6x10<sup>5</sup> | 15.99 | 1137392 | 6183 | 0.54 | 98.06 | 40.41 | 54.51 | 1131209 | Y | Y | Y |
|  | 2.6x10<sup>4</sup> | 18.99 | 1307167 | 4073 | 0.31 | 96.43 | 24.11 | 29.18 | 1303094 | N | Y | Y |
|  | 2.6x10<sup>3</sup> | 22.4 | 1452489 | 3391 | 0.23 | 91.19 | 21.16 | 46.71 | 1449098 | N | Y | Y |
|  | 2.6x10<sup>2</sup> | 25.34 | 1313739 | 827 | 0.06 | 67.8 | 4.91 | 10.02 | 1312912 | N | Y | Y |
|  | 26 | 29.34 | 779997 | 484 | 0.06 | 38.92 | 2.32 | 5.40 | 779513 | N | N | Y |
|  | 2.6 | 32.36 | 621358 | 8 | 0 | 2.42 | 0.04 | 0.25 | 621350 | N | N | N |
|  | 0 | 35.04 | 579159 | 8 | 0 | 0.47 | 0.01 | 0.20 | 579151 | N | N | N |
| **3** | 2.6x10<sup>6</sup> | 12.22 | 3760214 | 1780362 | 47.35 | 100 | 13486 | 15324 | 1979852 | Y | Y | Y |
| **SISPA** | 2.6x10<sup>5</sup> | 15.99 | 1318547 | 825545 | 62.61 | 100 | 5620 | 7583 | 493002 | Y | Y | Y |
|  | 2.6x10<sup>4</sup> | 18.99 | 1074343 | 502065 | 46.73 | 100 | 3716 | 4323 | 572278 | Y | Y | Y |
|  | 2.6x10<sup>3</sup> | 22.4 | 320287 | 86097 | 26.88 | 100 | 628 | 809 | 234190 | Y | Y | Y |
|  | 2.6x10<sup>2</sup> | 25.34 | 62371 | 6666 | 10.69 | 84.36 | 44.85 | 86.80 | 55705 | N | Y | Y |
|  | 26 | 29.34 | 54312 | 438 | 0.81 | 23.63 | 2.88 | 10.34 | 53874 | N | Y | Y |
|  | 2.6 | 32.36 | 28572 | 53 | 0.19 | 16.19 | 0.28 | 0.7 | 28519 | N | N | N |
|  | 0 | 35.04 | 1251455 | 177 | 0.01 | 19 | 1.00 | 2.12 | 1251278 | N | N | N |
| **4** | 2.6x10<sup>6</sup> | 12.22 | 880977 | 130315 | 14.79 | 100 | 835 | 1334 | 750662 | Y | Y | Y |
| **S-P** | 2.6x10<sup>5</sup> | 15.99 | 688161 | 109020 | 13.76 | 100 | 705 | 1020 | 579141 | Y | Y | Y |
|  | 2.6x10<sup>4</sup> | 18.99 | 1175085 | 105719 | 9.00 | 100 | 668 | 973 | 1069366 | Y | Y | Y |
|  | 2.6x10<sup>3</sup> | 22.4 | 80352 | 6901 | 8.59 | 100 | 43.84 | 58.94 | 73451 | N | Y | Y |
|  | 2.6x10<sup>2</sup> | 25.34 | 344436 | 9026 | 2.62 | 89.22 | 56.69 | 73.17 | 335410 | N | Y | Y |
|  | 26 | 29.34 | 105886 | 384 | 0.36 | 14.28 | 2.22 | 9.71 | 105502 | N | Y | Y |
|  | 2.6 | 32.36 | 151936 | 41 | 0.03 | 9.17 | 0.21 | 0.76 | 151895 | N | N | N |
|  | 0 | 35.04 | 734944 | 18 | 0 | 4.52 | 0 | 0.48 | 734926 | N | N | N |

\*Burrow-Wheeler Aligner

**Table 2.** Detailed statistics of SARS-CoV-2 genome assembly after SISPA amplification coupled with Illumina sequencing.

| Viral titre (pfu/ml) | Ct value | Gene | Reference assembly |  |  | *De novo* assembly |
| --- | --- | --- | --- | --- | --- | --- |
|  |  |  | The percentage (%) of genome covered | Average overage depth | SD |  |
| 2.6x10<sup>6</sup> | 12.22 | orf1ab | 100 | 21439.7 | 20844.3 |  |
|  |  | S | 100 | 2560.8 | 776.9 |  |
|  |  | orf3a | 100 | 3203.5 | 606 |  |
|  |  | E | 100 | 2630 | 446.3 |  |
|  |  | M | 100 | 3114.5 | 976.3 |  |
|  |  | orf6 | 100 | 4874 | 233.5 |  |
|  |  | orf7a | 100 | 8860.4 | 2037.3 |  |
|  |  | orf7b | 100 | 13001.8 | 298 |  |
|  |  | orf8 | 100 | 13765.3 | 821.8 |  |
|  |  | N | 100 | 19803.9 | 3615.6 |  |
|  |  | orf10 | 100 | 10446.2 | 2341.1 |  |
| Genome<br>(including non-coding regions) |  |  | 100 | 17104.6 | 19211.0 | Y |
| 2.6x10<sup>5</sup> | 15.99 | orf1ab | 100 | 9651.8 | 10201.6 |  |
|  |  | S | 100 | 990 | 584.9 |  |
|  |  | orf3a | 100 | 1070.9 | 420 |  |
|  |  | E | 100 | 690.3 | 258.5 |  |
|  |  | M | 100 | 391.5 | 162.5 |  |
|  |  | orf6 | 100 | 474.8 | 123.6 |  |
|  |  | orf7a | 100 | 2244.3 | 667.2 |  |
|  |  | orf7b | 100 | 3182.9 | 60.7 |  |
|  |  | orf8 | 100 | 2892.8 | 434.7 |  |
|  |  | N | 100 | 1831.2 | 434.2 |  |
|  |  | orf10 | 100 | 1991 | 351.1 |  |
| Genome<br>(including non-coding regions) |  |  | 100 | 7246.3 | 9438.9 | Y |
| 2.6x10<sup>4</sup> | 18.99 | orf1ab | 100 | 6192.5 | 5919.4 |  |
|  |  | S | 100 | 962.4 | 434.5 |  |
|  |  | orf3a | 100 | 1075.7 | 404.4 |  |
|  |  | E | 100 | 711.6 | 283.4 |  |
|  |  | M | 100 | 539.8 | 218.6 |  |
|  |  | orf6 | 100 | 460.1 | 112 |  |
|  |  | orf7a | 100 | 2256 | 788.2 |  |
|  |  | orf7b | 100 | 3455.9 | 34.8 |  |
|  |  | orf8 | 100 | 3153 | 308.7 |  |
|  |  | N | 100 | 2635.3 | 401.4 |  |
|  |  | orf10 | 100 | 1615.3 | 346.6 |  |
| Genome<br>(including non-coding regions) |  |  | 100 | 4812.3 | 5484.8 | Y |
| 2.6x10<sup>3</sup> | 22.4 | orf1ab | 100 | 1100 | 1099.9 |  |
|  |  | S | 100 | 133.7 | 113.5 |  |
|  |  | orf3a | 100 | 62 | 28.4 |  |
|  |  | E | 100 | 61 | 19.1 |  |
|  |  | M | 100 | 75.5 | 62.7 |  |
|  |  | orf6 | 100 | 76.1 | 24.3 |  |
|  |  | orf7a | 100 | 390.8 | 116.6 |  |
|  |  | orf7b | 100 | 423.1 | 24.4 |  |
|  |  | orf8 | 100 | 257.1 | 74.7 |  |
|  |  | N | 100 | 147.7 | 56.4 |  |
|  |  | orf10 | 100 | 153.3 | 12.4 |  |
| Genome<br>(including non-coding regions) |  |  | 100 | 826 | 1028.4 | Y |
| 2.6x10<sup>2</sup> | 25.34 | orf1ab | 95.5 | 78.3 | 123.7 |  |
|  |  | S | 84.5 | 12 | 9.6 |  |
|  |  | orf3a | 0 | 0.5 | 0.8 |  |
|  |  | E | 0 | 0.6 | 0.7 |  |
|  |  | M | 0 | 0 | 0 |  |
|  |  | orf6 | 0 | 0.4 | 0.6 |  |
|  |  | orf7a | 100 | 10.4 | 3.5 |  |
|  |  | orf7b | 100 | 6.8 | 1.1 |  |
|  |  | orf8 | 100 | 12.2 | 2.1 |  |
|  |  | N | 100 | 21.7 | 10.8 |  |
|  |  | orf10 | 100 | 25.6 | 3.9 |  |
| Genome<br>(including non-coding regions) |  |  | 89.6 | 59 | 109.1 | N |
|  | 29.34 | N/A |  |  |  | N |
|  | 32.34 | N/A |  |  |  | N |
|  | 35.04 | N/A |  |  |  | N |

SD, Standard deviation

### The SISPA (method 3) for WGS of SARS-CoV-2 is reproducible

We applied the SISPA method to four other cell cultured SARS-CoV-2 isolates (EDB-2, EDB-8, EDB-10 & EDB-12), to assess reproducibility of the method to give depth of coverage across the whole genome (Table 3, Figure 2). The SARS-CoV-2 sequencing reads distribution is shown in Figure 2 and resulted in full genome assembly for all four additional isolates. The percentage of viral reads obtained after sequencing that mapped to the reference SARS-CoV-2 genome resulting in complete genome assembly was between 33% to 84% for the SARS-CoV-2 viruses tested (Table 3). We obtained a high average coverage depth across the genome for all viral genes, the mean average being 46181.62 nucleotides per base (ranging from 16935.4 to 70780 nucleotides per base) (Table 3). The coverage depth per bp position for each viral gene was, at least 20,000 bp per base for orf1ab gene (from 20,330 to 88,920), 10,000 bp for orf7b (ranging from 12,817 to 29,776) and orf8 (from 11,888 to 27,124), 5,000 bp for orf7a (from 5,222 to 18,832), 3,000 for S gene (ranging from 3351 to 21,345 bp per base), and orf3a (ranging from 3,600 to 15,605), 2,000 for E and orf6 (from 2,155 to 16,381), and 1,500 for M gene (from 1,675 to 15,179). A very high depth of coverage was achieved for N (above 30,000 bp per base) and orf10 genes (above 85,000 bp per base) (Table 3).

**Figure 2.**
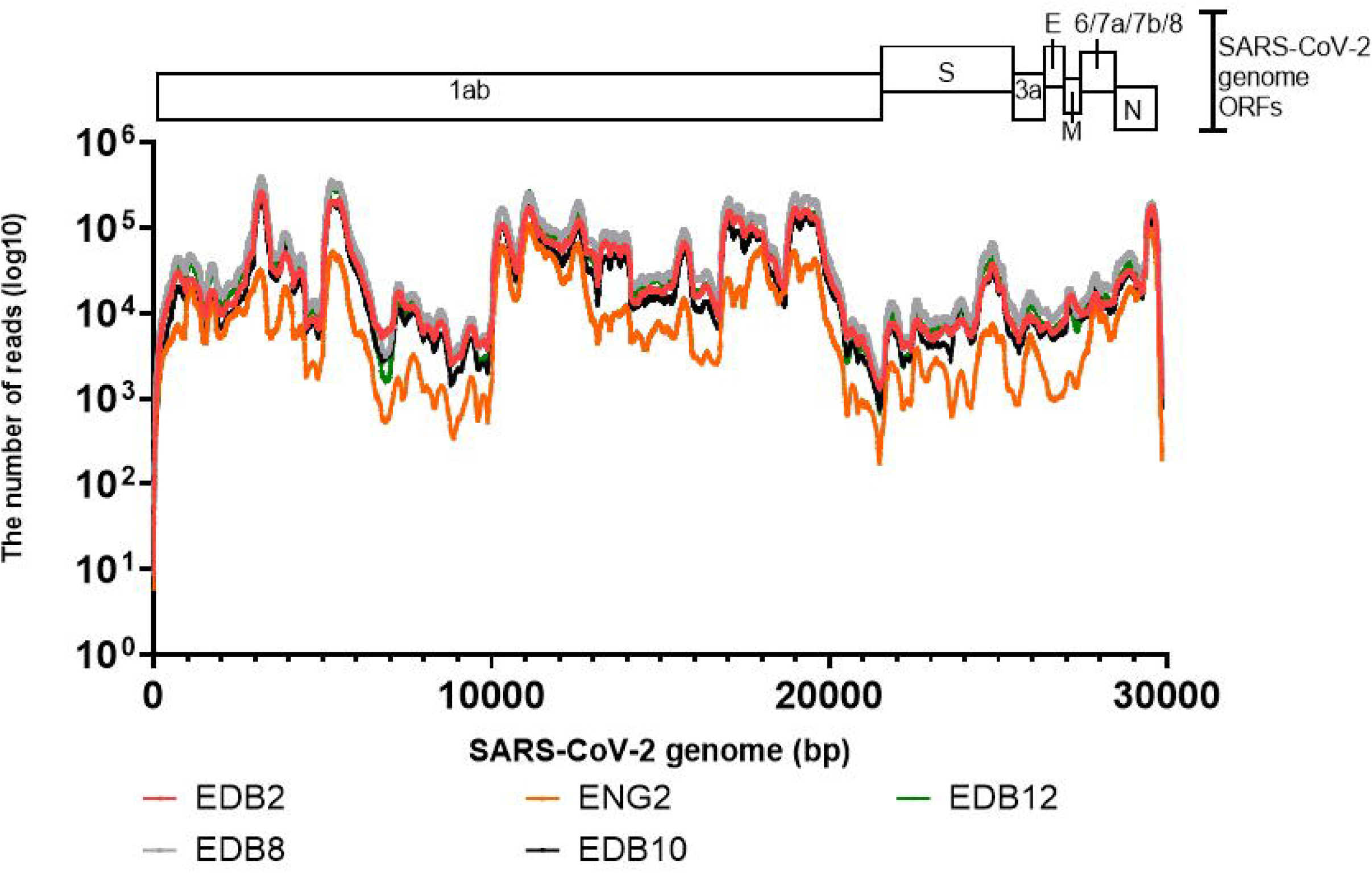
Read distributions aligning to SARS-CoV-2 viral genome after Sequence-Independent, Single-Primer Amplification (SISPA) coupled with Miseq Ilumina sequencing for whole-genome sequencing. SARS-CoV-2 virus strains used include hCov-19/England/02/2020 (Eng-2) and hCov-19/Scotland/EDB1827/2020 (EDB-2), hCov-19/Scotland/EDB2398/2020 (EDB-8), hCov-19/Scotland/EDB2057/2020 (EDB-10) and hCov-19/Scotland/EDB2405/2020 (EDB-12). Number of reads at each genome position is plotted.

**Table 3.** Reproducibility of genome coverage and average depth for the individual viral genes of SARS-CoV-2 viruses sequenced in this study. Sequence-Independent, Single-Primer Amplification (SISPA) technique was used to recover viral RNA followed by Miseq Illumina sequencing.

| SARS-CoV-2 isolate | Eng-2 (Titre) |  |  | EDB-2 (Titre) |  |  |
| --- | --- | --- | --- | --- | --- | --- |
| Genes | GC<sup>+</sup> (%) | DC<sup>\$</sup> / bp | SD<sup>^</sup> | GC<sup>+</sup> (%) | DC<sup>\$</sup> / bp | SD<sup>^</sup> |
| orf1ab | 100 | 20330 | 23576.5 | 100 | 60851.6 | 62374.8 |
| S | 100 | 3351.2 | 2153.9 | 100 | 12865.7 | 10103.3 |
| orf3a | 100 | 3600 | 1883 | 100 | 9480.8 | 1391.4 |
| E | 100 | 2510.6 | 565.5 | 100 | 8813.8 | 319.3 |
| M | 100 | 1675.5 | 394.6 | 100 | 11363.7 | 3911.1 |
| orf6 | 100 | 2155.2 | 144.9 | 100 | 16381.8 | 2029.6 |
| orf7a | 100 | 5222.4 | 2547.5 | 100 | 13544.9 | 1703.9 |
| orf7b | 100 | 12817.1 | 726.8 | 100 | 18326.3 | 356.3 |
| orf8 | 100 | 11888 | 1855.3 | 100 | 17113.4 | 1333.8 |
| N | 100 | 30319.9 | 27469.1 | 100 | 50962 | 47834.8 |
| orf10 | 100 | 85634.3 | 13447.4 | 100 | 175087.4 | 22253 |
| Genome |  | 16935.4 | 22570.4 |  | 48758.9 | 58082.7 |
| Number of reads mapped to reference |  | 1708791 (33%) |  | 4919631 (79%) |  |  |
| Total paired reads |  | 5156768 |  | 6209676 |  |  |

| SARS-CoV-2 isolate | EDB- 8 (Titre) |  |  | EDB -10(Titre) |  |  |
| --- | --- | --- | --- | --- | --- | --- |
| Genes | GC<sup>+</sup> (%) | DC<sup>\$</sup> / bp | SD<sup>^</sup> | GC<sup>+</sup> (%) | DC<sup>\$</sup> / bp | SD<sup>^</sup> |
| orf1ab | 100 | 88920.9 | 93125.5 | 100 | 48578.8 | 53822.8 |
| S | 100 | 21345.2 | 19076.8 | 100 | 10955.9 | 8806.2 |
| orf3a | 100 | 15605 | 3046.8 | 100 | 7486.4 | 1778.2 |
| E | 100 | 13196.2 | 768.9 | 100 | 7271.4 | 177 |
| M | 100 | 15179 | 3580.1 | 100 | 8397.3 | 2216 |
| orf6 | 100 | 15631.4 | 2166.9 | 100 | 12144 | 1032.1 |
| orf7a | 100 | 18832.7 | 4354.9 | 100 | 12510 | 2857.5 |
| orf7b | 100 | 29776.2 | 703.5 | 100 | 20697.5 | 541.7 |
| orf8 | 100 | 27124.4 | 2414.6 | 100 | 14315 | 21838 |
| N | 100 | 66978.3 | 48080.8 | 100 | 41626.3 | 36425.1 |
| orf10 | 100 | 188689 | 22483.6 | 100 | 128792.6 | 17961.1 |
| Genome |  | 70780 | 85008 |  | 39010.3 | 49216.3 |
| Number of reads mapped to reference |  | 7149251 (68%) |  | 3934982 (84%) |  |  |
| Total paired reads |  | 10389994 |  | 4676652 |  |  |

| SARS-CoV-2 isolate | EDB -13 (Titre) |  |  |
| --- | --- | --- | --- |
| Genes | GC<sup>+</sup> (%) | DC<sup>\$</sup> / bp | SD<sup>^</sup> |
| orf1ab | 100 | 71457.9 | 77597 |
| S | 100 | 15308.9 | 13398.7 |
| orf3a | 100 | 11414 | 2831.2 |
| E | 100 | 9169 | 910.3 |
| M | 100 | 10576.5 | 2415.9 |
| orf6 | 100 | 10509.1 | 1572.2 |
| orf7a | 100 | 15260 | 4550.5 |
| orf7b | 100 | 26180.5 | 571.7 |
| orf8 | 100 | 21716.8 | 3032.2 |
| N | 100 | 57886.3 | 53962 |
| orf10 | 100 | 194686 | 25369.3 |
| Genome |  | 55423.5 | 70955.1 |
| Number of reads mapped to reference |  | 5759620 (83%) |  |
| Total paired reads |  | 6879474 |  |

\* GC, genome coverage. \$ DC, depth of coverage per base pair. ^ SD, Standard deviation.

### Comparison of limits of detection for SARS-CoV-2 virus for the four amplification methods

To assess the limit of detection and the limits on full genome sequence assembly for each of the four different method protocols we used a ten-fold dilution series of the Eng-2 SARS-CoV-2 virus. As anticipated, for all methods the genome coverage and depth of coverage correlated with virus titre (Figure 3 and Suppl. Figure 1). Our results showed that a full genome sequence could be assembled with a low abundance of viral genetic material, minimum viral titre of 2.6×10^3^ pfu/ml (CT:22.4) using the SISPA or S-P protocols (methods 3 and 4) (Table 1, Figure 3, Suppl. Figure 1). The percentage of reads mapped to reference genome at this low virus titre was between 2% to 5% (S-P and SISPA, respectively). The average depth coverage for the SISPA method at 2.6×10^3^ pfu/ml (CT:22.4) virus load was 248 nucleotides per base (ranging from 1100 for orf1ab gene to 60 for arf3a and E genes) (Table 2 and Suppl. Figure 2). In comparison, the H-P and K-P protocols (methods 1 & 2) were able to produce full genome assemblies only when the input virus titre was high, above 2.6×10^6^ pfu/ ml (Table 1 and Figure 3). Even at this input however, the depth of coverage and percentage of mapped viral reads recovered with H-P and K-P methods was low (below 1%). For these reasons, these two methods were excluded from further analysis due to overwhelming competition with non-specific or host genome sequences that was not permissive for assembling the SARS-CoV-2 viral genome.

**Figure 3.**
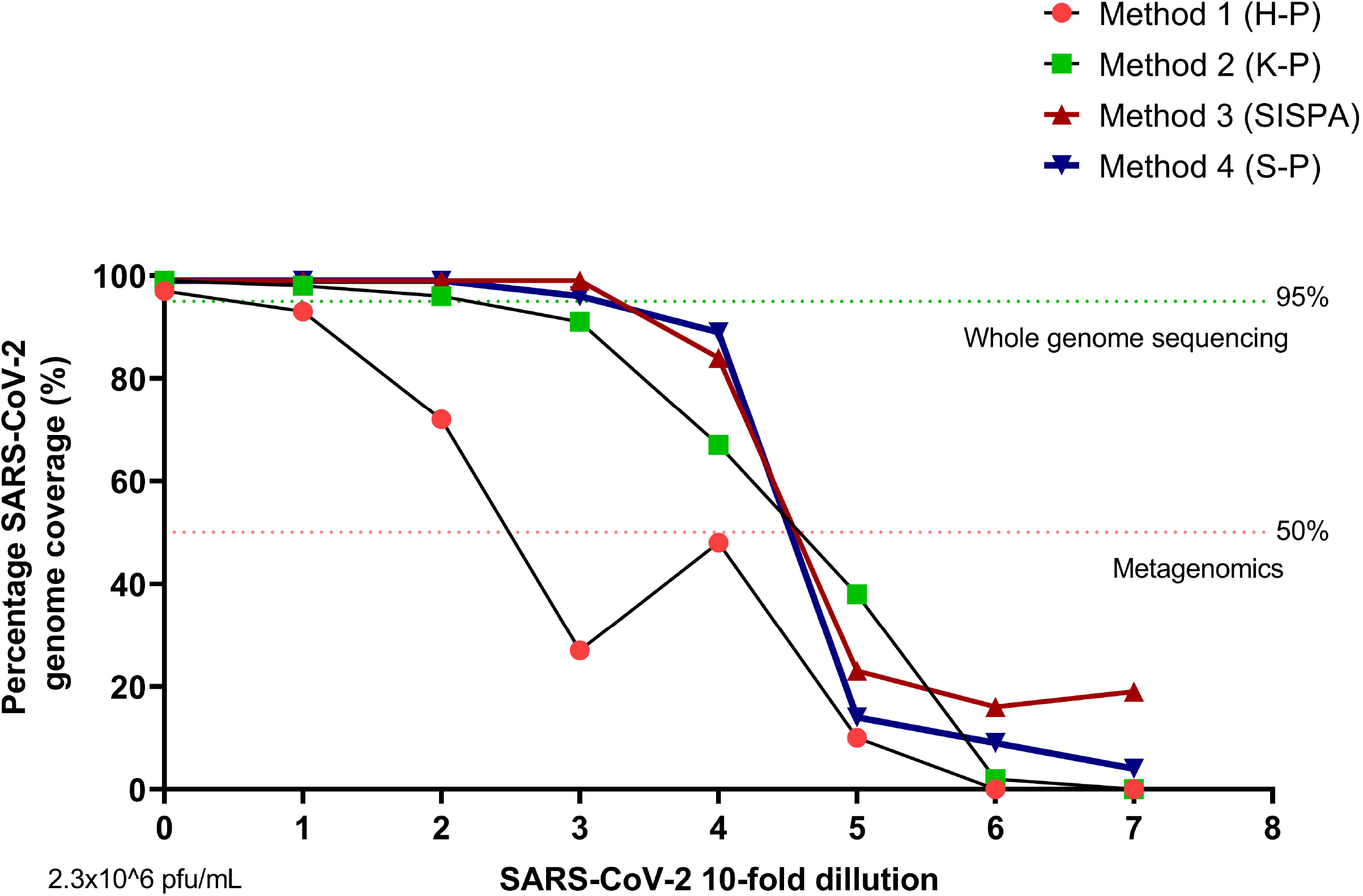
The percentage of SARS-CoV-2 genome coverage after applying the four different random priming methods coupled with next-generation sequencing. The SARS-CoV-2 virus was 10-fold serially diluted (from 2.3×106 pfu/mL, mark as “0” on x axis).

Below 10^3^ pfu/ml we were not able to assemble full or near full SARS-CoV-2 viral genome by any of the methods applied (Table 1, Figure 3 and Suppl. Figure 1). However, at 2.6 x 10^2^ pfu/ ml both the SISPA and S-P methods did give over 80% coverage of the SARS-CoV-2 genome (Table 1 and Figure 3). The in-depth analysis of the SISPA method indicated that the 84% of genome coverage resulted in 100% coverage of the following open reading frames; orf7a, orf7b, orf8, N and orf10, whilst orf1ab was 95.5% covered and S 84.5% covered (Table 2 and Suppl. Figure 2). The genome areas of orf3a, E, M and orf6, a contiguous region between nucleotides 25400 and 27350 of the SARS-CoV-2 genome had no coverage (Table 2 and Suppl. Figure 2).

The average depth of coverage and the number of reads mapped to the reference genomes using the SISPA or S-P method drastically decreased (to the level below 1%) below inputs of 10^2^ pfu/ ml of virus with no reproducible mapping possible at this level. This suggests that the limit for whole genome assembly of SARS-CoV-2 using SISPA method (or S-P method) is above 10^3^ pfu/ ml but depending on the area of genome of interest SISPA could give detail down to 10^2^ pfu/ml (Ct=25.34).

Although the genome coverage of virus with titre below 10^2^ pfu/ml (Ct=25.32) decreased drastically, it was still possible to detect SARS-CoV-2 genome after SISPA or S-P amplification using metagenomics. The limit of detection using metagenomics was 2.6×10^1^ pfu/ ml (CT: 29.34) using SISPA or S-P methods (Table 1). No virus was detected by metagenomics above CT value of 30 in this study.

### Full genome recovery of SARS-CoV-2 and A(H1N1)pdm09 influenza virus multiplexed in a single reaction

The SISPA method produced the most viral reads of any of the four methods employed at all the dilutions tested, therefore we used this method to recover full genome sequences of SARS-CoV-2 and A(H1N1)pdm09 influenza virus mixed together in single sample (Figure 4, panel B). To assess the limit of detection for full genome assembly of SARS-CoV-2 virus in a mixed viral sample, 10-fold diluted SARS-CoV-2 viral RNA (initial concentration 2.6×10^6^ pfu/ml and Ct value of 13.61) was spiked with a constant amount of A(H1N1)pdm09 viral RNA (Ct=24.88+/-0.19). The full genome sequence of A(H1N1)pdm09 and SARS-CoV-2 was assembled from each sample by *de novo* assembly and reference mapping (Figure 4, panel B). In all samples sequenced the influenza virus genome was fully sequenced by *de novo* methodology. The full genome of SARS-CoV-2 was assembled from the initial viral titre of 2.6×10^5^ pfu/ ml (CT:17) and above only (Figure 4B and Table 4). This differed to the scenario of SARS-CoV-2 alone when we were able to WGS the virus at a viral titre greater than 2.6×10^3^ pfu/ ml (CT value of 22.4) (Figure 4, Panel A and B). In the mixed viral samples a SARS-CoV-2 viral titre of 2.6×10^3^ pfu/ ml (CT: 23.51) allowed assembly of 70% of coronavirus genome sequence and full genome sequence of A(H1N1)pdm09 influenza virus (Figure 4, Panel B). The cumulative percentage of reads mapped to the reference viral genomes (SARS-CoV-2 and A(H1N1)pdm09) was between 37%-51% for the whole dilution series and a decrease in number of SARS-CoV-2 virus reads correlated with an increased number of A(H1N1)pdm09 virus reads (y= −0.6758x+35.468, R^2^=0.43), but not with an increase in reads of host *GalGal* genome (R^2^=0.0003) (Figure 4, panel B). For instance, at high SARS-CoV-2 viral load (CT: 13.61 and virus titre 2.6×10^6^ pfu/ ml), 38% of total sequencing reads mapped to the SARS-CoV-2 viral genome and 13% to the A(H1N1)pdm09 virus genome whereas at SARS-CoV-2 virus titre of 10^3^ pfu/ ml (CT:23.51), 8% of reads mapped to SARS-CoV-2 viral genome and 29% to H1N1 influenza virus genome (Figure 4, panel B). Importantly, the percentage of total “non-viral” unmapped reads (BWA-MEM unmapped neither to SARS-CoV-2 nor A(H1N1)pdm09) did not change and was the same for all the samples (52.63% +/- 6.96%) (Figure 4, panel B). H1N1 influenza virus stocks were produced in embryonated hens’ eggs, therefore the *Gallus gallus* (GalGal4.0) genome was used to map non-viral reads. The percentage of reads assembled to the host GalGal4.0 was similar in all samples (4-9% of total sequencing reads generated) in exception to the sample that contained high titres of both, SARS-CoV-2 and H1N1virus where only 1% of total reads were assembled to host GalGal4.0 reference genome (Figure 4, panel B).

**Figure 4.**
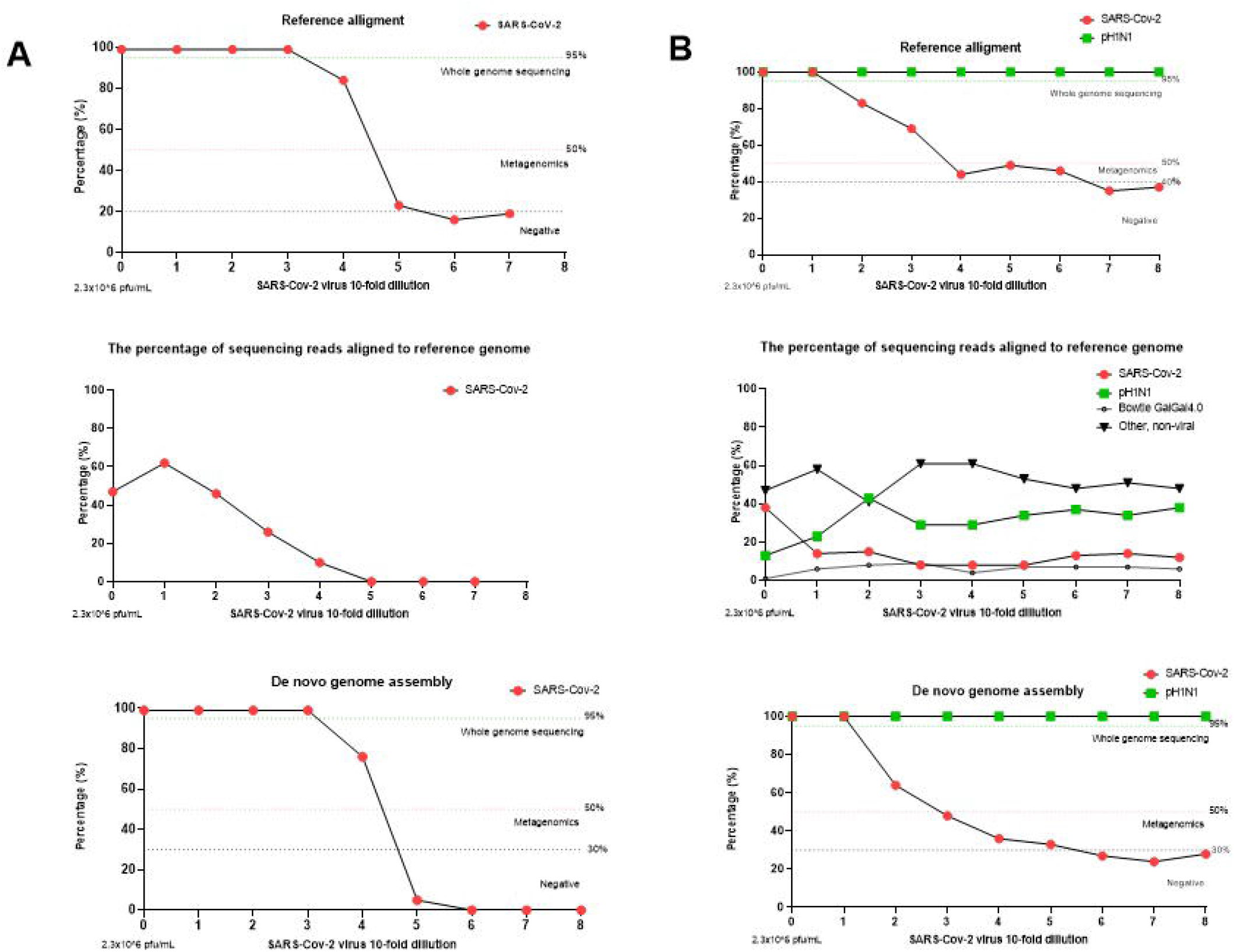
SARS-CoV-2 whole genome sequencing using Sequence-Independent, Single-Primer Amplification (SISPA) coupled with Miseq Ilumina sequencing. Left panel (A) represents single SARS-CoV-2 virus genome assembly (the percentage of genome coverage after reference mapping and de novo assemblies). The virus was 10-fold serially diluted, starting from viral load of 2.3 x 106 pfu/ml, mark as “0” on x-axis followed by 2.3 x 105 pfu/mL (mark as 1), 2.3 x 104 pfu/mL (mark as 2), etc. Right panel (B) represents genome assembly (the percentage of genome coverage after reference mapping and de novo assemblies) of two viruses, SARS-CoV-2 and influenza A(H1N1)pdm09 in mixed viruses single sample. SARS-CoV-2 virus was 10-fold serially diluted, starting from viral load of 2.3 x 106 pfu/mL, mark as “0” on x-axis, that was spiked with constant amount of H1N1 virus (7.4 x 106 pfu/mL).

**Table 4.** Genome assembly statistics of SARS-Cov-2 and pH1N1 influenza virus in single reaction. Sequence-Independent, Single-Primer Amplification (SISPA) technique was used to recover viral RNA followed by Miseq Illumina sequencing. 10-fold serially diluted SARS-CoV-2 virus was spiked with a constant amount of A/England/195/09 pH1N1 (7.4 × 10^6^ PFU/ml) pandemic influenza virus.

| SARS-CoV-2<br>Ct<br>value<br>(E-gene) | H1N1<br>Ct<br>value<br>(M-gene) | Total<br>number of<br>paired<br>reads | *De novo*<br>assembly (% of<br>genome<br>covered) |  | Reference<br>assembly (% of<br>genome covered) |  | Percentage of<br>reads mapped to<br>reference (%) |  | Metagenomics |  |
| --- | --- | --- | --- | --- | --- | --- | --- | --- | --- | --- |
|  |  |  | SARS-Cov-2 | H1N1 | SARS-CoV-2 | H1N1 | SARS-CoV-2 | H1N1 | SARS-CoV-2 | H1N1 |
| 13.61 | 24.76 | 1739986 | 100 | 100 | 100 | 100 | 38 | 13 | Y | Y |
| 16.97 | 24.64 | 489302 | 100 | 100 | 100 | 100 | 14 | 23 | Y | Y |
| 20.2 | 25.04 | 401355 | 65 | 100 | 83 | 100 | 15 | 43 | Y | Y |
| 23.51 | 24.85 | 1993925 | 48 | 100 | 69 | 100 | 8 | 29 | Y | Y |
| 26.5 | 24.84 | 754361 | 36 | 100 | 44 | 100 | 8 | 29 | Y | Y |
| 30.44 | 24.75 | 579777 | 33 | 100 | 53 | 100 | 8 | 34 | Y | Y |
| 33.24 | 25.23 | 450283 | 27 | 100 | 51 | 100 | 13 | 37 | Q | Y |
| 37.4 | 24.93 | 394865 | 24 | 100 | 35 | 100 | 14 | 34 | N | Y |
| 38.28 | 24.76 | 428735 | 28 | 100 | 37 | 100 | 12 | 38 | N | Y |

Q-questionable

## Discussion

Whole genome sequencing (WGS) is increasingly applied in clinical medicine as it has the potential to identify clinically actionable genetic variants informing early medical intervention. Importantly, WGS can act as valuable tool to monitor aberrations, such as mutations in viral genomes, following vaccination or anti-viral treatment, that could lead to therapy failures [40–45]. Recently, Wibmer et al. [44] have shown that SARS-CoV-2 variant 501Y.V2 (B.1.351), that contains two substitutions in S protein can escape from convalescent plasma neutralization antibodies (nMAbs) which can then result in lack of efficacy of S-based vaccines. Constant monitoring of SARS-CoV-2 viral genomes will be essential to control of SARS-CoV-2 virus spread, as antigenically distinct variants will reduce efficacy of spike-based vaccines available on the market globally. Furthermore, in addition to vaccination, new antiviral therapeutic agents to treat SARS-CoV-2 [46–49] by targeting viral genome regions such as *RdRP* polymerase gene are in development [50] which can lead to these areas of the genome changing and therefore affecting testing methods [51–53].

Although, both targeted and non-targeted WGS can generate whole genome sequences, the main advantage of non-targeted WGS is the absence of a prior assumption about the pathogens contain within the sample. The non-targeted identification of pathogens, therefore, allows the ability to detect any causative agent of infection (or disease outbreak), enables the identification of multiple pathogens in single reaction that could mask clinical output of disease, and circumvents issues caused by genetic variation in the genome that may affect targeted methods. In addition, information about the presence of bacterial species in samples and potential antibiotic resistance or virulent genes can also be recovered. Recent studies have shown that SARS-CoV-2 and other respiratory pathogens can co-exist in one host, causing respiratory infection [54–57]. Hence, it is of importance to examine for all potential pathogens in a sample as this might change the clinical output of disease and thus disease treatment.

In this study, we show a simple viral RNA template enrichment protocol coupled with Illumina sequencing multiplexed for whole genome sequencing of both SARS-CoV-2 and influenza A(H1N1)pdm09 viruses. This has been compared to the original protocol [32], with new changes including enhanced hexamer only and phi29 polymerase amplification. Presented in this study the SISPA protocol allowed for whole genome assembly of both viruses using only one primer in a sequence independent reaction. The percentage of SARS-CoV-2 viral reads obtained at high virus load ranged between 33% to 84% depending upon the sample and resulted in full coronavirus genome assemblies. However, percentage of reads ranging from 2% to 14% of viral-specific sequencing reads was enough to successfully assembly full or near full SARS-CoV-2 genome, depending upon status as either single or mix infection sample with influenza virus, respectively. Moreover, we have obtained high (13,486 nucleotides per base) average coverage depth for SARS-CoV-2 viral genome, with approximately 4489 nucleotides average coverage per base for S gene (ranging from 3351 to 21345.2 nucleotides per base) at high virus concentration that allows for polymorphism analysis of viral genome at 1% variant calling [39, 58]. Wolfel et al. [59] have shown that pharyngeal virus shedding is very high during the first week of symptoms, with a peak at 7.11 × 10^8^ RNA copies per throat swab on day 4, followed by an average titre of 3.44 × 10^5^ copies per swab after day 5 of infection whereas the average viral load in sputum samples was 7.00 × 10^6^ copies per ml, with a maximum of 2.35 × 10^9^ copies per ml at the same time point tested. Huang et al. [60] demonstrated that high virus titre of SARS-CoV-2 (Ct value around 15) can be still found in the sputum samples at one-week post-infection. The SISPA method presented here consequently, could be potentially applied on clinical samples received from symptomatic patients or critically ill patients, where high virus titre in the swab is expected [59–62] and allow for whole genome assembly of SARS-CoV-2 virus with high depth of coverage which would be useful for tracking any aberration in viral genome, eg. mutations following treatment or to monitor any secondary or co-existing infections.

For the whole genome assembly, we showed that the number of viral specific sequencing reads appear distributed between the RNA viruses contained in the sample rather than any host derived sequences and correlated directly with initial input viral load. Although the S-P technique presented in this study did not improve sequencing depth after Illumina sequencing, it might also be useful method to consider when Oxford Nanopore sequencing of SARS-CoV-2 is used as this method produces a longer dsDNA average fragment size of template for further library preparation (we obtained an average fragment size of 20Kb of dsDNA, ranging from 17.9 Kb to 22.2 Kb). An overwhelming number of non-viral sequencing reads were obtained after H-P or K-P methods resulting in lower than 1% of virus sequencing reads produced even when the high virus titre was used in this study. This should be considered when applying hexamer only based amplification as it can result in low depth of coverage, making it impossible to perform viral genome assembly when the virus titre is low and thus pre-detection methods are required so mapping can be directed rather than *de novo*. As compared to other recently published studies that utilize PCR-based targeted enrichment and either Illumina or Oxford Nanopore sequencing [10, 63, 64] the main advantage of SISPA (and/or S-P) method presented in this study is its simplicity (e.g. only one K-8N primer used), and possibility to apply the method to any unknown samples as no prior knowledge about pathogen is needed. As we showed here, this protocol was successfully applied to SARS-CoV-2 and influenza A(H1N1)pdm09 viruses mix infection in single reaction and allowed us to pull out whole genome sequences of both viruses. Interestingly, decreased number of SARS-CoV-2 viral specific sequencing reads loosely correlated with increased number of influenza A(H1N1)pdm09 virus specific (y= - 0.6758x+35.468, R^2^=0.43) but importantly we did not observe an increased in *GalGal* host genome sequencing reads, suggesting that the method presented here is capable of selectively recovering low abundance viral RNA genetic sequences. However, it is important to mention that in targeted whole genome sequencing where multiple pairs of primers are used, even though do it does not allow for assembly of multiple pathogens in singe reaction, the problem with generation of overwhelming number of host genome sequencing reads is also resolved. Hence, the sequencing method of choice depends on the aims of the study where the method is applied. ARCTIC network offers the most updated targeted whole genome sequencing methods (https://artic.network/ncov-2019).

Furthermore, we assessed the feasibility of virus identification and estimated its limit of detection for diagnosis of covid-19 infection or co-infection with influenza viruses. We showed that by using the SISPA or S-P protocols presented in this study, the full genome sequence can be assembled when initial viral titres are as low as 2.6×10^3^ pfu/ml for single SARS-CoV-2 virus in the sample and approximately 10^5^ pfu/ml viral titre (SISPA method) if it is a mixed infection of both viruses with influenza virus being at high titre. However, it is unknown how likely both viruses might be found at a high viral load in a single clinical sample or how one virus will influence the replication of another [65, 66]. We also assessed the detection limit for the amplification methods presented in this study using metagenomics approaches. The in-house metagenomics pipeline (Figure 5) enabled us to detect SARS-CoV-2 virus in the sample when the initial virus titre was approximately Ct value of 30 regardless of single or mix infection sample and no prior sequence information was needed. This might suggest that the method presented here should allow to detect asymptomatic or pre-symptomatic patients as median Ct value (for two genetic targets: the N1 and N2 viral nucleocapsid protein) reported by Arons et al. [67] for asymptomatic residents, pre-symptomatic residents, residents with atypical symptoms and residents with typical symptoms, were 25.5, 23.1, 24.2, and 24.8, respectively. Similar Ct value for asymptomatic and pre-symptomatic SARS-CoV-2 in Washington, US, were also showed by Kimball et al. [68]. Smith et al. [69] determined that the limit of detection with 100% detection for Abbott RealTime SARS-CoV-2 Emergency Use Authorization (EUA) is 100 copies/ml (n=80), with Ct mean and standard deviation was 26.06±1.03. Voges et al. [52] have compare the most common SARS-CoV-2 qRT-PCR assays developed by the China Center for Disease Control (China CDC), United States CDC (US CDC), Charité Institute of Virology, Universitätsmedizin Berlin (Charité), and Hong Kong University (HKU) and found that the most sensitive primer-probe sets are E-Sarbeco (Charité), HKU-ORF1 (HKU), HKU-N (HKU), CCDC-N (China CDC), 2019-nCoV_N1 (US CDC), and 2019-nCoV_N3 (US CDC), could partially detect SARS-CoV-2 at 1 (25%) and 10 (25-50%) virus copies per μL of RNA. Although the direct comparison between qRT-PCR and whole genome shotgun metagenomics is difficult to perform, as PCR-based methods targeting short fragment of genetic material and therefore aim only at detection which makes theirs limit of detection usually being low, eg Ct of 36 (ORF1 SARS-CoV-2 qRT-PCR) and 37 (E-gene qRT-PCR) [70] as compare to presented here SISPA-NGS along with our in-house metagenomics pipeline that lies at Ct of 30 (based on E-gene), shotgun metagenomics nevertheless deliver satisfactory results which should allow to detect symptomatic or asymptomatic covid-19 infected patients. Buchan et al. [70] have shown that among 1,213 specimens tested as SARS-CoV-2 positive, the median Ct values of covid-19 samples were 25.02 and 25.93 for ORF1 and E-gene, respectively, which indicate that the distribution of Ct values observed in symptomatic patients is approximately 5 Ct value above our metagenomics pipeline limit of detection. Metagenomics analysis of samples that contain less than Ct of 30 might be possible, however for that purpose a pre-processing step in the sample preparation might need to be applied such as DNase treatment, or viral concentrations techniques [71] that could potentially improve the efficacy of viral amplification and sequencing. Notably, the method presented here does not rely on primer specificity as compared to conventional qRT-PCR [72] and therefore any changes in viral genomes (mutations or deletion) do not impact the pathogen detection. Previous studies have shown active genetic recombination events in SARS-CoV-2 genomes which may reduce the accuracy of conventional qRT-PCR detection and thus the primers should be precisely chosen to address these challenges [31, 73–75].

**Figure 5.**
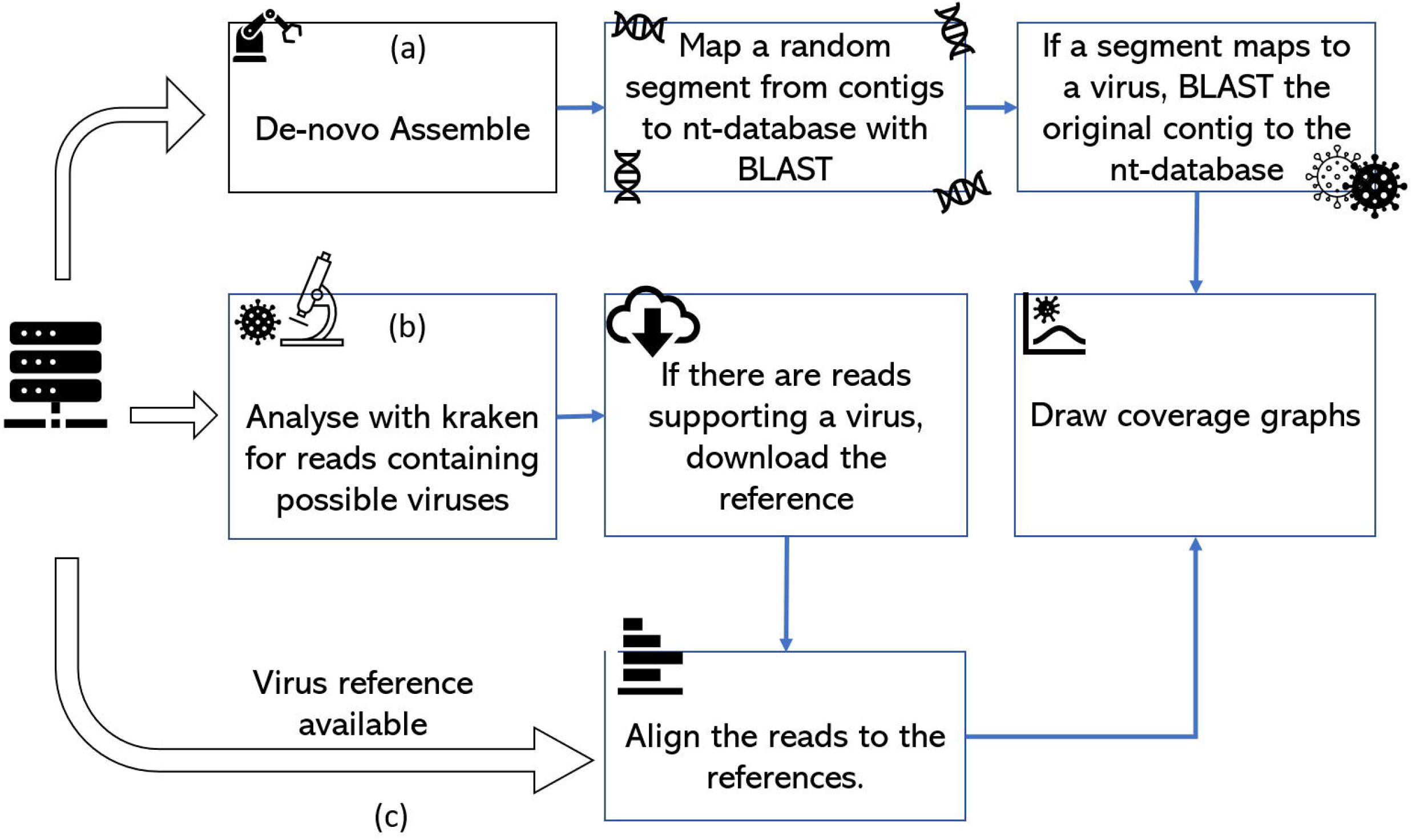
The diagram illustrates the analysis pipelines for the virus detection process. (a) High concentration viruses allow de novo assembly to construct the genome. The contigs are tested with BLAST and the nt-database for known viruses and the selected virus genomes coverage plotted. (b) The k-mer content of the reads is inspected with Kraken and if the presence of a virus is supported by a high number of reads, the references are downloaded and reads aligned to them. (c) If a reference genome is available, the reads can be directly aligned. For flows (b) and (c) the read depth against genome position can be plotted.

## Conclusion

In conclusion, the performance of four different random priming amplification methods to recover RNA viral genetic material (SARS-CoV-2) were compared in this study. The SISPA technique allowed for whole genome assembly of SARS-CoV-2 and influenza A(H1N1)pdm09 in mixed viruses single samples. We assessed limit of detection and characterization of SARS-CoV-2 virus which lies at 10^3^ pfu/ml (Ct, 22.4) for full-length SARS-CoV-2 virus genome assembly and Ct of 30 for virus detection. We also presented S-P technique that might be useful to apply for Oxford Nanopore real-time sequencing as no non-targeted primer-based protocol is available yet. The whole genome sequences recovered after applying SISPA (or S-P) method presented in this study are free of primer bias and allowed for polymorphism analysis. This method is predominantly useful for obtaining genome sequences from RNA viruses or investigating complex clinical samples (such as mixed infections in single reaction) as no prior sequence information is needed. The method might be useful to monitor SARS-CoV-2 virus changes such as mutation or deletions in virus genome, to perform simple and fast metagenomics detection and to assess general picture of diffrent microbes within the sample that might be useful to identify the other co-factors that correspond to covid-19 infection.

## Abbreviations

COG-UK: Covid-19 genomics UK consortium
Ct: Cycle threshold
FBS: Foetal bovine serum
*GalGal*: *Gallus Gallus*
Kb: Kilo bases
MDA: Multiple displacement amplification
NGS: Next generation sequencing
nMabs: Neutralising monoclonal antibodies.
Pfu: Plaque forming unit
RT: Reverse transcriptase
SISPA: Sequence independent single primer amplification
WGA: Whole genome amplification
WGS: Whole genome sequencing
ZIKA: Zika virus

## Declarations

### Ethics approval and consent to participate

Not applicable

### Consent for publication

Not applicable

### Availability of data and materials

The datasets used and/or analysed during the current study are available from the corresponding author on reasonable request.

### Author contributions

The work was conceptualized by KC and HS. Experimental work was executed by KC, CT, DB, GF and JF. The manuscript was written by KC and HS and edited by all authors.

## Acknowledgments

We acknowledge the Pirbright High Throughput Sequencing unit and provision of SARS-CoV-2 strains from Public Health England and Dr Christine Tait-Burkard at The Roslin Institute, UK.

