## supplementary figures for "A random priming amplification method for whole genome sequencing of SARS-CoV-2 and H1N1 influenza A virus"

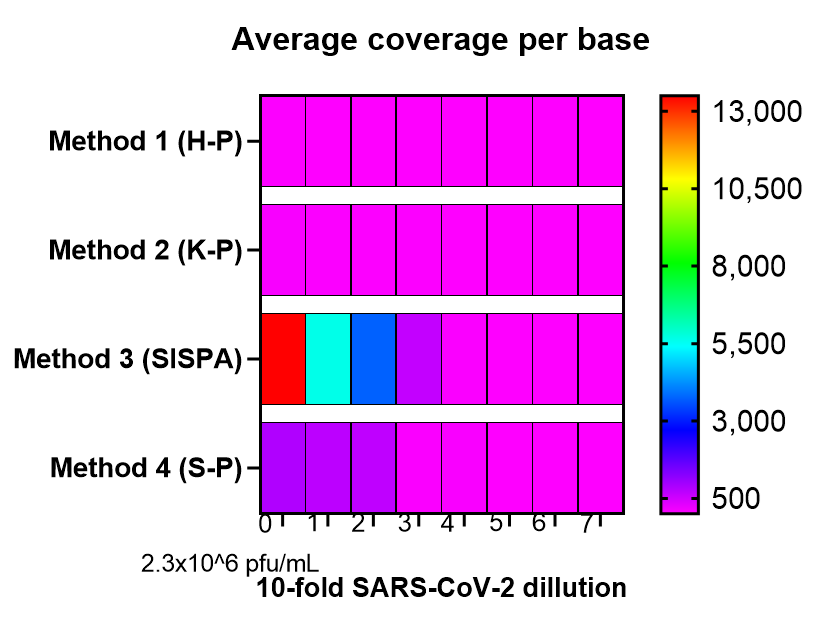

Suppl. Figure 1. The average depth of coverage obtained after SARS-CoV-2 sequencing using the four different random priming methods coupled with next-generation sequencing. The virus was 10-fold serially diluted (from 2.3 x10^6 pfu/mL, mark as “0” on x axis).

Suppl. Figure 2. Read distributions aligning to SARS-CoV-2 viral genome following Sequence-Independent, Single-Primer Amplification (SISPA) coupled with Miseq Ilumina sequencing. SARS-CoV-2 virus was 10-fold serially diluted, starting from viral load of 2.3 x 10^6^ pfu/mL (S0), 2.3 x 10^5^ pfu/mL (S1), 2.3 x 10^4^ pfu/mL (S2), etc. Number of reads at each genome position is plotted.
